# Hosts, parasites, and their interactions respond to climatic variables

**DOI:** 10.1101/079780

**Authors:** Timothée Poisot, Cynthia Guéveneux-Julien, Marie-Josée Fortin, Dominique Gravel, Pierre Legendre

**Affiliations:** Département de Sciences Biologiques, Université de Montréal; Québec Centre for Biodiversity Sciences; Department of Ecology and Evolutionary Biology, University of Toronto; Département de Biologie, Université de Sherbrooke

**Keywords:** beta diversity, species interaction networks, climatic niches, species distribution models, rodents, fleas

## Abstract

Aim: Although there is a vast body of literature on the causes of variation in species composition in ecological communities, less effort has been invested in understanding how interactions between these species vary. Since interactions are crucial to the structure and functioning of ecological communities, we need to develop a better understanding of their spatial distribution. Here, we investigate whether species interactions vary more in response to different climate variables, than individual species do. Location: Eurasia. Time period: 2000s. Major taxa: Animalia. Methods: We used a measure of Local Contribution to Beta-Diversity to evaluate the compositional uniqueness of 51 host–parasite communities of rodents and their ectoparasitic fleas across Eurasia, using publicly available data. We measured uniqueness based on the species composition, and based on potential and realized biotic interactions (here, host-parasite interactions). Results: We show that species interactions vary more, across space, than species do. In particular, we show that species interactions respond to some climatic variables that have no effect on species distributions or dissimilarity. Main conclusions: Species interactions capture some degree of variation which is not apparent when looking at species occurrences only. In this system, this appeared as hosts and parasites interacting in different ways as a reponse to different environments, especially the temperature and dryness. We discuss the implications of this finding for the amount of information that should be considered when measuring community dissimilarity.

---

Ecological communities are made of species, but also of the interactions among them. Understanding how community structure changes requires addressing the variability of these two components. Biogeography has developed a very strong corpus of theory and empirical evidence pertaining to the fact that species occur at different locations because of random chance, favourable habitat (environmental filtering), and the ability of these species to establish the biotic interactions they require to persist. Yet the spatial distribution of species interactions has been comparatively less looked at. Poisot *et al.* (2015) suggested that species interactions, at any given location, can vary because of neutral effects (species abundance), functional effects (species traits), or the effect of a suite of other unspecified factors. Interactions are known to react in stochastic ways to population abundances (Canard *et al.,* 2014), and are therefore inherently more variable. One of the possible factors that has not been evaluated is the effect of environmental conditions on the existence of interactions. If interactions respond to the environment, then ecological communities will vary across space. Here we investigate whether knowledge of environmental conditions helps to explain species interaction variability across space.

One way to investigate the variability of ecological communities is through the use of spatially replicated systems: by describing the presence/absence of species originating from the same regional pool at different localities and documenting their interactions, it is possible to compare these communities to gain insights about why and how ecological communities vary. Because this is done within a shared regional species pool, this approach reveals the role of other factors than species compositional changes on the variation of community structure. Information about both species and their interactions can be efficiently represented using ecological networks, and the recent years saw the development of approaches to quantify the variation of these networks (Poisot *et al.,* 2012). In parallel, advances in the quantification of *β*-diversity allow the identification of hotspots of variation, *i.e.* localities that through their unique composition, have a high contribution to the overall *β*-diversity (Legendre & De Cáceres, 2013; Legendre, 2014; Legendre & Gauthier, 2014). In this manuscript, we bring these families of approaches together, describing the variation of community structure across space, and identifying the mechanisms and environmental variables responsible for this variation.

Understanding species within their communities, and by extension communities themselves, can be done through a quantification of the species’ Grinnellian and Eltonian niches (Devictor *et al.,* 2010). To summarize, species contribute to community structure first by being present within the community (which assumes that the local environmental conditions are amenable to their persistence), and second by fulfilling a functional role within this community, which is in part defined by the way they interact with other species (Coux *et al.,* 2016). Communities therefore differ it two complementary ways: first, because they harbour different species; second because whichever species they share might interact in different ways (Poisot *et al.,* 2015). This dissimilarity in both species composition and species interactions, over space and time, has raised increasing empirical attention in recent years (Carstensen *et al.,* 2014; Vizentin-Bugoni *et al.,* 2014; Olito & Fox, 2015; Trøjelsgaard *et al.,* 2015). Although the drivers of species distributions have been well elucidated in the past decades, there is virtually no knowledge (nor hypotheses) regarding how species interactions should be distributed in space. Most of the earliest datasets on ecological interactions replicated across space (*e.g.* Havens, 1992) assume that interactions are constant: two species will always interact if they are found in the same location, environment, etc. Because empirical data (most of which stems from host-parasite interactions, but also from mutualistic systems) contradict this assumption, there is an urgent need to revise our understanding of how community structure should be defined at both the local and global scale (Trøjelsgaard & Olesen, 2016). Specifically, we need to do so in a way that accounts simultaneously for the variability of species occurrences, and for the variability of species interactions.

Building on the framework put forth by Poisot *et al.* (2015), it is possible to develop quantitative hypotheses regarding the variation of species presences, interactions, and the relationship between them. An interaction between any pair of species is only possible when the two species of this pair co-occur. Therefore, because two species may co-occur but not interact (whereas the opposite is impossible), the dissimilarity of interactions is greater than or exactly equal to the dissimilarity of species composition; it cannot be smaller if the two dissimilarities are measured from the same community data (Poisot *et al.,* 2012). There are two important consequences to this fact. First, species composition across multiple localities should be less dissimilar than species interactions. Second, interaction dissimilarity will produce a clearer picture of how communities differ, because the distribution of interactions is intrinsically more variable than that of species: even when two species are found together, they may not interact locally. Most importantly, species are nested within interactions (at least from a purely mathematical standpoint), whereby nodes/species are embedded into edges/interactions – therefore, describing interactions is a more informative way of depicting community structure, which not only includes but actually supersedes the usual approach of communities-as-lists-of-species.

We will test whether species interactions respond to environmental conditions, using interaction data on rodents and fleas in Eurasia (Hadfield *et al.,* 2013). Host-parasite systems offer an exceptional natural laboratory to investigate these questions. First, parasites require an interaction with their hosts to persist, so that we except a strong matching in the distribution of parasites and their hosts (Poulin *et al.,* 2011). Second, the effect of the parasite on its hosts fitness is not strong enough to trigger large mortality events (Krasnov *et al.,* 2002), so we do not anticipate repulsion of parasites and hosts in space. Third, fleas being macro ecto-parasites, they experience the same environment as their hosts, and therefore measuring their response to the same environmental conditions is justified. Finally, this rodents-fleas system has a very broad spatial distribution in Eurasia, covering a range of climatic regions. Because of all of these characteristics, this is a perfectly suitable system in which we can test the effect of environmental conditions on species interactions in space.

This study demonstrates how, within a biogeographic perspective, species interactions reveal another dimension in which ecological communities vary; importantly, species interactions not only capture the variability of species distributions, but also capture another layer of environmental conditions. In particular, we show that (i) species interactions are more variable, and allow the identification of more sites with unique contributions to *β*-diversity, than species only; and (ii), species interactions react to climatic variables of their own, in addition to capturing most of the climatic variables acting on species distributions. We discuss these results in the light of our current definition of ecological communities, and highlight ways of refining this definition in order to make more accurate and realistic predictions about community structure.

## MATERIALS AND METHODS

We measure how species composition and species interactions in host-parasite communities vary across environmental gradients. Specifically, we use novel methods to quantify the compositional uniqueness of localities based on different definitions of community structure (Legendre & De Cáceres, 2013), then identify the climatic variables involved in driving the dissimilarity between localities. An interactive document allowing to reproduce all analyses (including downloading all data from their source) is available as supplementary information (Appendix S2 in supporting information); it requires the knitr package to be compiled from R into HTML or into the PDF presented as Appendix S1 in supporting information.

### Species interaction data

The species interaction data were taken from Hadfield *et al.* (2013). They describe species interactions between rodents (121 species) and ecto-parasites (206 species of fleas) at 51 locations throughout Eurasia. This system is species-rich, and likely originates from successive co-speciation events within pairs of interacting species (Hadfield *et al.,* 2014). These data were collected by combing a large number of individual rodents, then identifying the fleas that were collected (Krasnov *et al.,* 2004). The data were downloaded from the *mangal* database [Poisot *et al.* (2016a); http://mangal.io/data/dataset/4/]. The communities (where “community” is defined as the species and interactions detected at one location) have between 3 and 27 (median 11) hosts, 7 and 40 (median 19) parasites, and 12 and 226 (median 63) interactions between them. Out of 326 species, 94 were observed only once, and 43 were observed at more than 10 locations.

For every location, we define two levels of analysis. First, the *realized* interactions; this corresponds to interactions that where reported to occur within individual locations in the original data. Second, the *potential* interactions; this corresponds to the interactions that could happen given the information contained in the entire dataset. For example, if parasite *P_j_* and host *H_k_* do not interact locally, but interact in at least one other location, there will be an interaction between them in the *potential* interaction network. These two levels encapsulate hypotheses about the filtering of species and interactions: *potential interactions* are what is expected if species vary in their spatial distribution, but interactions are entirely fixed; conversely, *realized interactions* are the interactions observed at each location, and therefore account for the spatial variability of interactions in addition to that of species (Poisot *et al.,* 2012). For a given location, the *realized* and *potential* interaction networks have an equal number of species, but the *realized* network has as many or fewer interactions than the *potential* one. The extent to which the *realized* and *potential* interactions differ is measured using *β'*_OS_ (Poisot *et al.,* 2012); in brief, this measure is, for every community, the pairwise dissimilarity between **R** and **Q**. Values close to 0 mean that all *potential* interactions are *realized* (weak interaction filtering), whereas values close to unity indicate that most potential interactions have been filtered out (strong interaction filtering).

This distinction between realized and potential interactions, as we detail below, is of great importance. There are many causes to the variation of species interactions, ranging from the distribution of functional traits (Olesen *et al.,* 2011), expressed at macro-ecological scales, to neutral and random-chance events (Canard *et al.,* 2014), expressed at the micro-ecological scale. Rodents in particular are known to change part of their phenology in response to climate (Bozinovic & Rosenmann, 1989; Aars & Ims, 2002), and this in turn can affect the reproductive success of parasites. In short, we argue that potential interactions are more likely to reflect the evolutionary history of the species pairs (this is evidenced in this system by the strong phylogenetic signal in species interactions – Hadfield *et al.* (2014)), while the realized interactions are more likely to reflect how this species pairs reacts to a set of local environmental conditions; specifically, changes in local conditions may, through their impact on species or on the interaction directly, prevent potential interactions from being realized. From a species distribution point of view, potential interactions offer the possibility to look at the co-distribution of interacting pairs of species. Indeed, any effect of environmental variables on the spatial distribution of potential interactions is highly suggestive of the fact that species that interact also distribute nonrandomly. Local factors affecting species interactions can blur whatever signal exists due to species co-distribution: this requires to estimate this effect through the statistical signal of the models describing the effect of the environmental variables on both realized and potential interactions.

### Climatic variables data

We downloaded the 19 *BioCLIM* data (Hijmans *et al.,* 2005) at a geographic resolution of 5 arc minutes. The data for each location were then extracted using the GPS coordinates of the sampling location. Since the precise spatial extent that was sampled around each location is unknown, we deemed more conservative to use a relatively coarse spatial resolution to capture the general environmental conditions around each site. Preliminary analyses revealed that the results are qualitatively unchanged when using different resolutions for the climatic data.

### Quantification of species and interactions variation

Our approach is represented in Figure 1. We use the method put forth by Legendre & De Cáceres (2013), where the overall *β*-diversity between sites of a spatially replicated sampling is measured as the variance of a community data matrix **Y**, and noted *β*_**Y**_. **Y** is a binary matrix with locations as rows and items on which to measure the dissimilarity as columns. These matrices are defined such that *Y(l, i),* that is the value in row *l* and column *i* of matrix **Y**, is 1 if item *i* is found at location l, and 0 otherwise. For site by species matrices, the row sums give the richness at the locations, and the column sums give the number of occurrences of the species. We define four community data matrices, which are special cases of **Y**. **H** has host species in columns, and **P** has parasite species in columns. These first two matrices will generate a baseline estimate for the dissimilarity of localities based on species composition. Finally, we also define **R**, with *realized* interactions in columns, and **Q** with *potential* interactions in columns.

**Figure 1.**
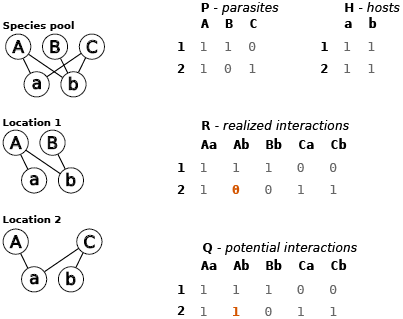
Illustration of the construction of the four community data matrices. The networks observed at every location are sampled from a regional pool of species (hosts a, b; parasites A, B, C), pictured at the top left. The two locations vary because they do not have the same species, nor do the shared species interact in the same way. The four matrices corresponding to this simplified situation are on the right side. Note that the interaction between host b and parasite A is 1 in the matrix of potential interactions for location 2; we know from the regional pool that it can happen, even though it was not observed at this location.

As per the recommendations in Legendre & Gallagher (2001), we applied the Hellinger transformation to the **Y** matrices. The dissimilarity matrices (**D**_**H**_, **D_P_**, **D_R_**, and **D_Q_**) were computed using the Euclidean distance on transformed data (Legendre & De Cáceres, 2013). The transformed matrices were then analysed to measure the Local Contributions to Beta-Diversity (LCBD). LCBD is a quantification of how much every location (row of the **Y** matrix) contributes to the overall dissimilarity, presented as a vector **l_Y_**. We interpret this value as a measure of the *originality* of a location: a large contribution to beta-diversity indicates that the location has a set of species or interactions that is different from the overall species pool. The values of LCBD are tested for statistical significance under a permutation scheme, specifically by re-distributing species or interactions across locations. This tests whether LCBD values are larger than expected from random variations in species composition (**H**, **P**) or species interactions (**Q**, **R**). We assume that LCBDs larger than expected by chance indicate that the locality is *unique* with regard to the species or interactions found therein. We used the default significance threshold of *α* = 0.05 (after Holm-Bonferroni correction), with 9999 random permutations of each column of the four community data matrices.

To summarize, for each community data matrix **Y** representing hosts, parasites, potential, and realized interactions, we measure the *β*-diversity (*β*_**Y**_), the matrix of pairwise distance between sites (**D_Y_**), the extent to which each location contributes to *β*_**Y**_ (**l_Y_**), and the statistical significance of each element *l*_**Y**_(*i*).

### Ordination and variable selection

We investigated the extent to which the dissimilarity among communities was driven by climatic variables using RDA (see *Appendix S1* in supporting information; Legendre & Legendre, 2012). The 19 *bioClim* variables were put in a matrix **E** with one location per row. To identify the most significant climatic variables, we used stepwise model building using forward variable selection and AIC-like statistics over 9999 replicate runs for each matrices. This approach yields four models of the form RDA(**Y** ~ **E**) (that is, we explain the structure of the **Y** matrix using the environmental conditions at each site as predictors), for which we extract **e_Y_**, *i.e.* a vector of significant climatic variables to explain the structure of **Y**. In each model, we also record the rank of each climatic variable; variables selected early have a stronger contribution to the structure of community composition.

### Interpretation

Taking a step back from the statistical analysis, we anticipate three possible outcomes with regard to the effect of environmental conditions on species and species interactions dissimilarity. In the first situation, environmental conditions have an impact on species, but not on their interactions: this would suggest that interactions are fully predicted by species traits or properties that are independent from environmental conditions (or that the suite of bioclimatic variables used did not contain the relevant information). In the second situation, species and interactions react to the same set of environmental predictors: this would suggest that species and interactions share environmental drivers, and that interactions covary very strongly with species co-distribution. Finally, the alternative situation is one where interactions and species have some unique predictors: this would indicate that interactions are responding to aspects of the local environment that have no linear effect on species.

The difference in how realized and potential interactions respond to the environment will allow us to delineate the role of co-distribution versus local drivers of interactions. Specifically, unless the realized interactions are affected by drivers that have no impact on potential interactions, we will conclude that most of the variation in interactions over space is tied to species co-distribution rather than local variation driven by the environment. If drivers are shared between realized and potential interactions, especially if they are not shared with species, these environmental variables affect interactions at both scales.

## RESULTS

### Species and interactions vary across space

In Figure 2, we show that the 51 locations have realized interactions that are not the same as those found at the regional level – if interactions where perfectly conserved from the *potential* to *realized* steps, all locations would have *β_OS_ =* 0 (no dissimilarity). Although the strength of interactions varies across space, *β'*_OS_ measures whether their *presence* varies too, *i.e.* they are realized at one location, but not at another. Specifically, the mode of the distribution of *β'*_OS_ is around 0.3, indicating that the filtering of interactions from the regional pool to local communities is comparable to other rodents–parasite communities (Poisot *et al.,* 2013 reported values between 0.2 and 0.4 in a central European system). Using the Legendre & De Cáceres (2013) approach reveals that interactions vary *more* than species. Specifically, *β***_R_** ≈ 0.94, *β***_Q_** ≈ 0.90, *β*_**H**_ ≈ 0.80, and *β*_**P**_ ≈ 0.79. That the variation in **R** is larger than that in **Q** is expected, as the *realized* interactions represent the action of additional ecological mechanisms involved in the filtering of *potential* interactions. That localities in this dataset exhibit different interactions, and have varying strengths of filtering from the potential to the realized interactions, raises the question of identifying which of these localities have the strongest contribution to *β*-diversity.

**Figure 2.**
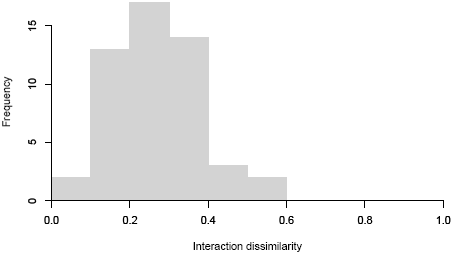
Distribution of *β'*_OS_ across the entire dataset. Local communities tend to keep a majority of the regional interactions, as evidenced by dissmilarity values going down to 0, with nevertheless some variability.

### Species and interactions contribute differently to site uniqueness

Using the LCBD approach, we measure the extent to which each location contributes to the beta-diversity of hosts, parasites, local, and potential interactions. These results are presented in Figure 3. For each distance matrix, we clustered the locations using partition around the medoids (Kaufman & Rousseeuw, 1990), and selecting the number of medoids that gave the smallest silhouette. This yielded respectively three clusters for the hosts (east; north west; south west), and two for the parasites; that hosts and parasites have different clusters suggest that they do not perfectly co-distribute. The same three clusters are found for the potential interactions, and the realized interactions (north; south); this is expected since interactions are constrained by species composition, and so the finer division of host communities is also observed when using interactions as the proxy for community structure.

**Figure 3.**
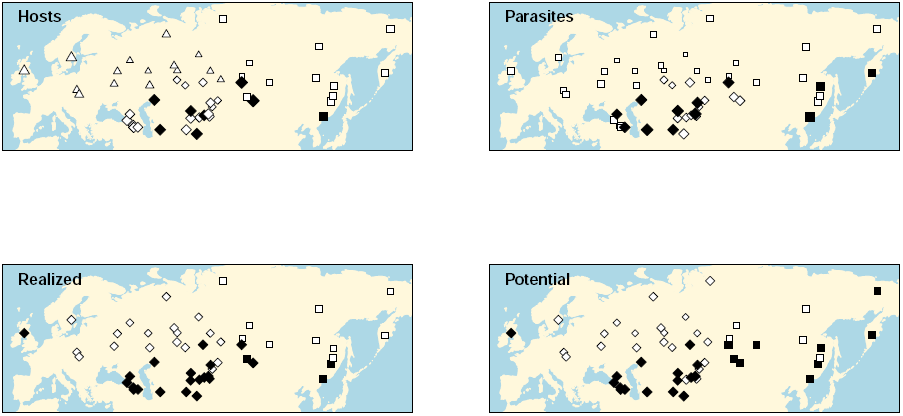
The beta-diversity matrices were divided in clusters (indicated by different symbols). Black-filled symbols have significant Local Contribution to Beta Diversity (LCBD) values. The size of each symbol is proportional to its LCBD value. Note that there is no correspondance between the symbols used to denote cluster identity across the four panels.

Overwhelmingly, the locations with significant contributions to beta-diversity are located in the southernmost half of the dataset, in more desertic areas (*i.e.* Turkestan, Taklamakan, and Gobi deserts). These locations have environmental conditions that vary from what is found in the remainder of the dataset (see next section). There are 8 host locations with significant contributions to beta diversity and 12 for parasites, *i.e.* locations that have unique species compositions. In contrast, there are 23 locations with unique realized interactions and 25 for potential interactions. With six exceptions, realized and potential interactions agree on which locations are unique.

### Beta-diversity of species and interactions is structured by the environment

In (Figure 4, we present the results of db-RDA analyses of the four beta-diversity matrices, using the *bioClim* variables as predictors (additional informations such as number of constrained axes and inertia are given in the Supplementary Material). As in (Figure 3 (and because environmental variables tend to be spatially autocorrelated), locations from the same cluster, and locations with significant contributions to beta-diversity, tend to occupy the same area of the space defined by the canonical axes. This suggests two things. First, clustering of communities as a function of their species or interaction composition does indeed reflect that some species/interactions assemblages are specific to some combinations of environmental conditions. Second, locations with significant contributions to beta-diversity are unique because their local environmental conditions select unique species, and within this local species pool promote unique interaction assemblages.

**Figure 4.**
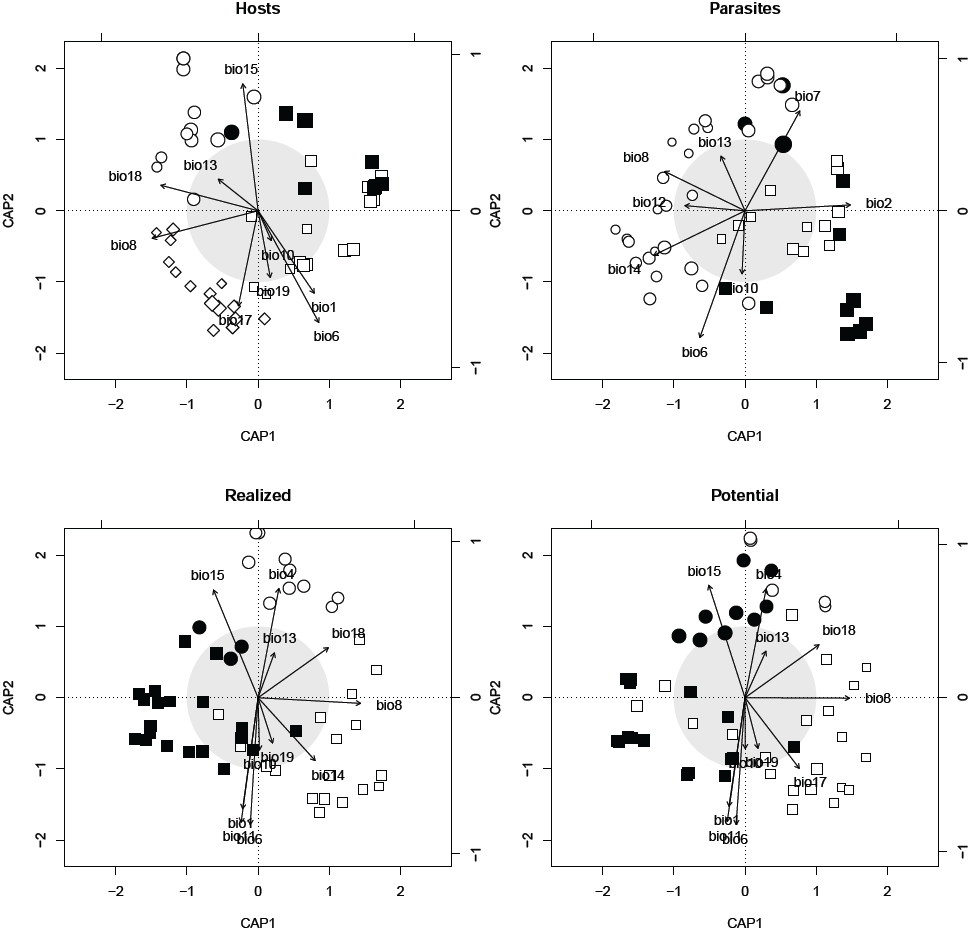
Ordinations of the four distance matrices, based on the scaled bioclimatic variables. Shape represents the cluster to which each location belongs, symbol size scales with LCBD, and filled symbols have significant LCBD. The ordination diagrams where produced using the capscale function in *vegan,* hence the axes are labelled CAP1 and CAP2.

Finally, in Table 1, we present the variables that have been retained during stepwise model building. The most important variables are the minimum temperature during the coldest month (`bio06`), and the minimum temperature during the wettest and coldest quarters (`bio08` and `bio10`). Some precipitation-related variables were associated with hosts and interactions, but not parasites. Meanwhile, a number of variables where associated with parasites only (and, surprisingly, not to interactions). Finally, two variables (`bio04`, seasonality of temperature; `bio11`, mean temperature of warmest quarter) where associated with interactions only. From Figure 4, it appears that the locations with the most significant unique assemblages (in all matrices) also have low values of temperature during the wettest quarter (`bio08`), whereas locations with non-significant assemblages have high values of this variable.

**Table 1.** List of bioclim variables retained during the model selection (also used in Figure 4). Numbers identify the rank in the forward-selected model. Variables 4 and 11 (in bold) are unique to interactions. Hosts, **H**; parasites, **P**; realized interactions, **R**; potential interactions, **Q**.

| bioclim variable | Explanation | H | P | R | Q |
| --- | --- | --- | --- | --- | --- |
| 06 | Min temp. cold. month | 1 | 1 | 1 | 1 |
| 08 | Mean temp. of wettest quarter | 2 | 3 | 2 | 2 |
| 10 | Mean temp. of coldest quarter | 5 | 4 | 6 | 5 |
| 13 | Precipitation of wettest month | 4 | 5 | 9 | 4 |
| 01 | Annual mean temp. | 3 |  | 3 | 3 |
| 18 | Precipitation of warmest quarter | 6 |  | 5 | 6 |
| 15 | Precip. seasonality | 7 |  | 4 | 8 |
| 19 | Precipitation of coldest quarter | 8 |  | 10 | 7 |
| 17 | Precipitation of driest quarter | 9 |  |  | 9 |
| 02 | Mean diurnal range |  | 2 |  |  |
| 05 | Max temp. warm. month |  | 6 |  |  |
| 12 | Annual precip. |  | 8 |  |  |
| 14 | Precipitation of wettest month |  | 7 |  |  |
| **04** | Temp. seasonality |  |  | 7 | 10 |
| **11** | Mean temp. of warmest quarter |  |  | 8 | 11 |
| 03 | Isothermality |  |  |  |  |
| 07 | Temp. ann. range |  |  |  |  |
| 09 | Mean temp. of driest quarter |  |  |  |  |
| 16 | Precipitation of wettest quarter |  |  |  |  |

## DISCUSSION

The *β* diversity of interactions was higher than the *β*-diversity of species. Although notable, this is not a surprising result for at least two reasons. First, in any community there are many more interactions than species, and therefore interactions capture more of the variation of community structure than species do. Second, interactions are probabilistic events (Poisot *et al.,* 2016b) that can vary even if the two species involved are consistently co-occurring. Here, we show that part of this variability happens independently of species traits or attributes, and is instead driven by local environmental conditions. Since the information in species presence/absence is by definition included in the information about species interactions, it stands to reason that we would more adequately describe community structure and variation by systematically considering species interactions. This is emphasized by the fact that measuring LCBD indices based on interactions revealed more (over twice as many) unique sites than measuring the LCBD of species (Figure 3). Quantifying the *β*-diversity of communities as based only on their species composition (i) consistently underestimates *β*-diversity and (ii) consistently underestimates the uniqueness of localities in the dataset. A species-centered vision of diversity, in short, does artificially homogenize the structure of communities by missing important sources of variation.

An additional result of our study is that, although species and their interactions in this system do share a lot of their bioclimatic predictors, interactions (**R** and **Q**) responded to two variables that had no bearing on species (**P** and **H**) composition (Table 1). We recognize that this can, in part, stem from the mismatch of scales between the observation of species interactions and the observation of climatic variables; in itself, this is an intriguing question: what is the scale at which ecological interactions are affected by the environment? Methods such as Moran Eigenvector Maps (Legendre & Gauthier, 2014) can help pinpoint the right scale at which each environmental variable is acting on the system – yet they are not applicable here, as they would require the measurement of the environmental conditions that happen directly where and when interactions are sampled, and not to be extracted from interpolated bioclimatic data. Nevertheless, our results highlight a challenge for current efforts to add biotic interactions to species distribution models (Boulangeat *et al.,* 2012; Araújo & Rozenfeld, 2014; Blois *et al.,* 2014): from additional predictors, species interactions become variables that must first be predicted themselves based on local environmental conditions. This is in addition to interactions requiring first and foremost the presence of both species (Gravel *et* al., 2011) to be realized. Nevertheless, this problem may prove less complicated based on the observation that there is little difference (besides their relative importance) in the predictors of potential and realized interactions. What this suggests is that climatic variables act on the distribution of *species pairs that can interact,* and the signal at the level of realized interactions is *inherited* from the co-distribution of potential interaction partners. Or in other words, the signal of potential interactions is a predictor of the co-occurrence of interacting species. As suggested in Poisot *et al.* (2015), interactions are filtered *after* the two species involved have co-distributed – it therefore stands to reason that environmental conditions that do not affect the co-distribution of the two species can affect the realization of the interaction locally.

The results we present here echo previous empirical evidence on rodent-parasite interactions. In particular, we report that communities close to desertic areas tend to have unique interactions, more so than they have unique species assemblages. Rosenzweig & Winakur (1969) identified micro-habitat features, like soil’s resistance to sheer stress that, through their effect on vegetation cover, modified the rules for competition between rodents species in deserts. Similarly, Kotler & Brown (1988) suggested that because deserts are not as resource-rich environments as the usual rodents habitats, many rules of foraging are modified. This includes increased among-patch mobility by hosts, and the overlap between diets of co-occurring species – these two facts have the potential to modify the rules of intra and inter-specific transmission of ectoparasites thereby creating unique communities in deserts. Kotler (1984) also identified an increase in predation risk in deserts, which are open habitats: this too changes the way rodents behave, and notably the moments of the day where they are active. The joint constraints of finding food and avoiding predation can very well have cascading effects on the way hosts are exposed to parasites and the way parasites are transmitted. More recently, Bordes *et al.* (2015) examined the idea that habitat fragmentation (which in rodents was modelled by the intensity of deforestation, reducing the amount of trees with a dense canopy in favour of more open shrubs and grasses) can change the host-ectoparasite interaction networks. They found that increasingly open environments had increasingly different network structures (without looking into the detail of how interactions are organized within these networks). There is, in effect, a series of empirical evidence pointing to the fact that rodent-parasite interactions can be affected by the environment. Therefore, although our results should lead to consider interaction networks as inherently dynamical objects that respond to the environment around them, they are not entirely surprising from an ecological point of view.

Across the entire spatial extent of the dataset, variables associated with interactions only capture warm temperatures (`bioll`) and temperature seasonality (`bio04`); this is an interesting contrast to the most important variables for all matrices, that are associated with colder temperatures – this may suggest that interactions respond to modifications of host phenology when exposed to warmer temperatures, despite this aspect of the environment having no impact on host/parasite species occurrence. Krasnov *et al.* (2004) previously reported that temperature and precipitation had an impact on the specificity of fleas in this system. Our present study greatly expands this result: not only does it pick up the contributions of variables to species distribution in addition to species interactions, our approach works at the scale of each interaction (as opposed to the specificity, which aggregates across all interactions for a parasite). We are therefore confident that this method will allow researchers to study into greater detail, for a variety of systems, the way interactions respond to the environment.

The results of this study highlight two important notions that we hope will be debated by biogeographers. First, interactions contain intrinsically more information than species; second, interactions react to climatic variables in ways that seem to have no bearing on the concerned species. Taken together with the fact that the information on species occurrence is by definition nested within the information on species interactions, this points to the idea that we should be cautious when defining what a “community” is. In particular, we show that while describing the presence and absence of species at different locations is important, it misses a lot of information by systematically under-estimating the variability *and* the uniqueness of these locations. For these reasons, we believe that the more desirable way of representing community structure is to describe, not only the species, but also their interactions. Although this demands a much larger sampling effort, it is the correct approach not to neglect an entire aspect of what is community *structure:* species, not being independent entities, are organized in non-random ways through their interactions. Species interactions do also capture some information on species traits (Bartomeus *et al.,* 2016; Coux *et al.,* 2016), and adding traits as a key component of our definition of community is indeed another obvious next step (McGill *et al.,* 2006). As recent contributions expanded the framework of community dissimilarity to account for species traits (Laliberté & Legendre, 2010; De Cáceres *et al.,* 2013), and the present contribution shows that same work is possible with interactions, we hope that the method presented here is a solid enough methodological basis to start investigating whether for any system, traits or interactions hold the most relevant information.

## Supporting information

Appendix S1 – Supplementary material

Appendix S2 – Document and code source to generate the supplementary material

## Acknowledgements and contributions

TP, DG, PL and MJF designed the study; PL suggested how to compute LCBD from interaction matrices; CGJ and TP performed the analyses and contributed equally to this study; TP and DG wrote the first draft of the manuscript. All authors provided comments and helped with the interpretation. TP, DG, PL and MJF were funded by a NSERC Discovery Grant; TP was also funded by a FRQNT New Investigator award. DG and TP acknowledge financial support from the Canadian Institute for Ecology and Evolution.

## Biosketch

Timothée Poisot is a quantitative and computational ecologist at the Université de Montréal. His work focuses on the spatial and temporal variability of community structure. He specializes in the re-analysis of existing data, and the large scale aggregation of open data for ecological synthesis.

